# Multidimensional transcriptomics provides detailed information about immune cell distribution and identity in HER2+ breast tumors

**DOI:** 10.1101/358937

**Authors:** Fredrik Salmén, Sanja Vickovic, Ludvig Larsson, Linnea Stenbeck, Johan Vallon-Christersson, Anna Ehinger, Jari Häkkinen, Åke Borg, Jonas Frisén, Patrik L Ståhl, Joakim Lundeberg

**Affiliations:** Science for Life Laboratory, Division of Gene Technology, KTH Royal Institute of Technology, Stockholm, Sweden; Hubrecht Institute-KNAW (Royal Netherlands Academy of Arts and Sciences) and University Medical Center Utrecht, Cancer Genomics Netherlands, Utrecht, the Netherlands; Division of Oncology and Pathology, Department of Clinical Sciences, Lund University, Lund, Sweden; Department of Pathology and Cytology, Blekinge County Hospital, Karlskrona, Sweden; Department of Cell and Molecular Biology, Karolinska Institutet, Stockholm, Sweden; Equal contribution

## Abstract

The comprehensive analysis of tumor tissue heterogeneity is crucial for determining specific disease states and establishing suitable treatment regimes. Here, we analyze tumor tissue sections from ten patients diagnosed with HER2+ breast cancer. We obtain and analyze multidimensional, genome-wide transcriptomics data to resolve spatial immune cell distribution and identity within the tissue sections. Furthermore, we determine the extent of immune cell infiltration in different regions of the tumor tissue, including invasive cancer regions. We combine cross-sectioning and computational alignment to build three-dimensional images of the transcriptional landscape of the tumor and its microenvironment. The three-dimensional data clearly demonstrates the heterogeneous nature of tumor-immune interactions and reveal interpatient differences in immune cell infiltration patterns. Our study shows the potential for an improved stratification and description of the tumor-immune interplay, which is likely to be essential in treatment decisions.

## Introduction

The recent rise in global life expectancy is accompanied by increasing prevalence of age-related diseases such as cancer^1,2^. Our current understanding of cancer as well as the efficacy of therapeutic interventions is largely driven by information gained from cancer tissue. A particularly important aspect is related to the identities and distribution of immune cells within tumor tissue^3–5^. Current standard analyses mainly rely on morphology-based examination of individual tissue sections by pathologists combined with additional insight from immunohistochemistry using antibodies towards known marker proteins^6^. However, a discrepancy between clinical IHC-based and transcriptomics-based analyses has been noted in the classification of HER2+ breast cancers^7^. A number of models for immune response in cancer have been proposed and it is currently under debate whether a high level of lymphocyte infiltration in Basal-like and HER2+ breast cancers is connected to better prognosis in early stage cancers^8–10^. Two important types of lymphocytes associated with breast cancer are cytotoxic T-cells (CD8+) and helper T-cells (CD4+). Both of these cell-types are present at low amounts in non-malignant breast tissue but show large variation in malignant breast cancer tissue^11^. The infiltrating lymphocytes can be present in the stroma adjacent to the tumor area or directly infiltrate the tumor area and interact with the cancer cells, i.e. tumor infiltrating T-cells. The presence of tumor infiltrating CD8+ T-cells facilitates anti-tumor immunity and is thus correlated with better overall patient survival^12–14^. In contrast, the presence of intra-tumor CD4+ T-cells is negatively related to patient outcome^15^. The dense concentration of tumor-associated macrophages has also been found to correlate with negative overall survival and the occurrence of metastases^16–18^. Several additional types of immune cells have been detected in breast cancer tissue, but their exact functions in cancer development remain unknown^19^. The composition and function of tumor infiltrating immune cells can directly influence the efficacy of certain therapeutic approaches. Recently, cancer immunotherapy, in which immune cells are stimulated to target the tumor, has begun to show promising results^20,21^. In such cases, comprehensive cancer-immune cell interaction analyses prior to specific activation might prove pivotal to therapeutic success.

The standard methods for describing the immune cell composition of tumor tissue depend on predetermined histological and molecular immune markers and are limited regarding multiplexing applications. They also heavily rely on microscopic assessments by trained pathologists, but traditionally lack three-dimensional resolution. To overcome these obstacles, we designed a workflow that leverages computational analysis of Spatial Transcriptomics data for the functional and immune cell type analysis of HER2+ breast cancer tissue samples. The computational approaches used in previous reports^22,23^, which are solely based on gene expression values, are more suitable for tissue samples with clearly defined molecular structures. However, tumor samples are usually highly complex in terms of cell mixture and distribution; therefore, other approaches are needed to reliably describe the molecular properties of such samples. In the presented study, we use an approach called Latent Dirchlet Allocation (LDA)^24^. The method was originally developed to describe the distribution of topics within a text. Any given sample has a specific proportion of each topic, allowing each sample to contain several topics simultaneously. Recently, this method has been successfully applied to biological experiments, for example, in the analysis of RNA-seq data^25,26^. Since each spatial spot contains a mixture of several different cell-types, this method is well suited for Spatial Transcriptomics^22^ data and can be exploited to identify underlying gene expression structures. Spatial spots with similar proportions of the topics (herein called gene expression topics) are considered to contain similar cell mixtures and are pooled to gain more depth in the data. Here, we show that this approach can be used to classify tissue regions based on function rather than morphology. By combining Spatial Transcriptomics with robust computational tools, we can now present immune cell distribution with richer spatial information. The further integration of cross-sectioning with computational alignment enables us to generate 3D-images of global expression patterns and immune score distributions throughout the samples.

## Results

Three tissue sections were collected from each of the ten patients (A-J) diagnosed with HER2+ breast cancer included in this study (Supplementary Table 1). The collected sections were then subjected to Spatial Transcriptomics^22,23,27,28^ using circular spatial spots with a diameter of 100µm. Each tumor was carefully examined by a trained pathologist, who annotated the different morphological regions in each tissue section. These annotations either served as a reference for subsequent unsupervised analysis or were used directly to pick spatial spots from regions of interest. During unsupervised analysis, we established biological functions and determined immune cell composition at the transcriptional level by using LDA to identify underlying structures of gene expression patterns. The proportions of gene expression topics were further used for hierarchical clustering, the object of which was to organize spatial spots with similar gene expression patterns into distinct clusters. Expression data from spatial spots within each cluster were further collapsed to increase gene detection. Next, pathway analysis^29^ was carried out between clusters in order to detect spatial correlation between different regions across the tissue. We then applied xCell^30^, a method for describing immune cell composition, to gain spatial information about the complex tumor-immune cell landscape by determining the abundance of sixteen different immune cell-types. The method overview is presented in Fig. 1a.

**Figure 1.**
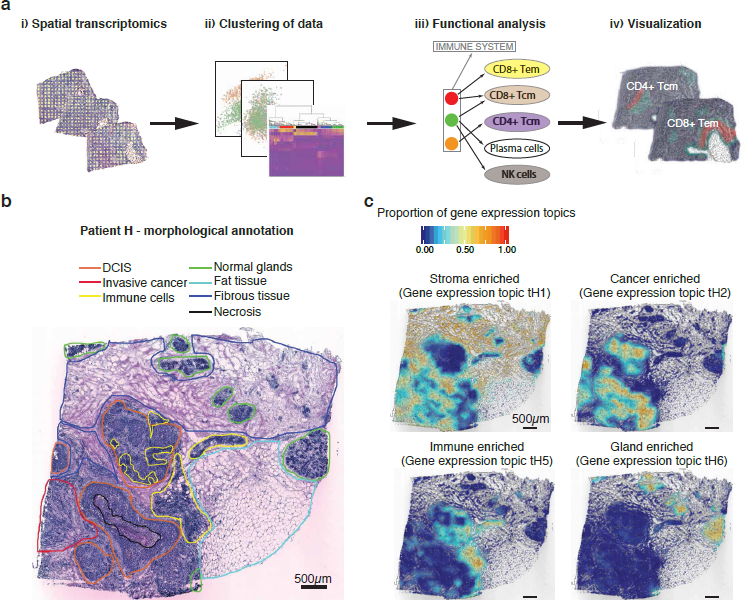
Overview of the presented approach. (a) i) The Spatial Transcriptomics method was applied to sections from ten patients (three sections from each biopsy); ii) spatial spots covering various regions of the tissue were selected and clustered; iii) spatial spots within each cluster were collapsed prior to pathway analysis and immune cell-type determination (Tem, effector memory T-cells; Tcm, central memory T-cells); iv) the data were visualized by superimposing tissue types on their spatial positions in the tissue. (b) Morphological regions were characterized by a pathologist and annotated into seven distinct categories: DCIS (orange); invasive cancer (red); immune cells (yellow); normal glands (green); fat tissue (cyan); fibrous tissue (blue); and necrosis (black). The black bar in the bottom left corner represents 500µm. (c) Interpolated tissue images that illustrate how four of the gene expression topics have clear morphological patterns. The color scale shows the proportion of each gene expression topic in specific tissue regions. Gene expression topic tH1 was prevalent in the stroma (both fibrous tissue and tumor stroma), tH2 was prevalent in cancer tissue (both DCIS and invasive), tH5 was associated with immune cells while tH6 was associated with normal glands. The black bar in the bottom left corner of each tissue section represents 500µm.

The morphological examination of tumor sections revealed inter- and intra-patient tissue heterogeneity. Tumors from all ten patients had invasive components, but only tumors from four patients (A, G, H and J) contained regions annotated as ductal carcinoma *in situ* (DCIS). Tumors from six of the patients (E, F, G, H, I and J) contained regions with a high abundance of immune cells. Other major tissue types identified included fat tissue, fibrous tissue and normal glands (Fig. 1b, Supplementary Fig. 1a). To study gene expression differences and similarities between tumors, we generated triplicate transcriptomics data sets (*in silico* bulk) for each patient. Dimensionality reduction by Principal Component Analysis (PCA) revealed several separate groups of patients, indicating large interpatient variation (Supplementary Fig. 1b).

To further analyze the Spatial Transcriptomics data, we applied LDA to each patient sample. A number of gene expression topics, each described by a certain set of genes, was used in the analysis (Supplementary Table 2). The gene expression sampling was based on the circular spatial spots. We visualized the data in a way that is easy for the human eye to interpret by generating interpolated images of the spatial spot positions, values of gene expression topic proportions and binary images of tissue morphology (Supplementary Fig. 2). We projected the interpolated values across the tissue sections to display the spatial distribution of gene expression topics. A comparison with the pathologists’ annotations (Fig. 1b, Supplementary Fig. 1a) revealed that certain gene expression topics clearly overlapped with regions described to be stromal, cancer, immune and gland-enriched (Fig. 1c, Supplementary Fig. 3). This finding demonstrates that applying LDA to Spatial Transcriptomics data can provide valuable information about the tumor microenvironment.

Next, we clustered spatial spots based on the proportions of gene expression topics (Fig. 2a, Supplementary Fig. 4). To validate the clustering, we also visualized the proportions of gene expression topics in t-SNE space (Fig. 2b, Supplementary Fig. 5) and overlaid them with the tissue morphology (Fig. 2c, Supplementary Fig. 6a). The separation of spatial spots in t-SNE space corresponded well to the gene expression topic clusters, while the spatial cluster distribution over the tumor section closely resembled the manual annotation. Taken together, these results demonstrate that clustering spatial spots based on gene expression topic proportions is a reliable approach for characterizing the tumor microenvironment in samples. To analyze variation across the triplicate data sets of each patient, we calculated the proportions of spatial spots assigned to each cluster in each triplicate. Most clusters showed similar distributions across the three data sets with the small variations between sections most likely representing molecular variation in different parts of the tumor (Supplementary Fig. 6b). Interestingly, the clustering revealed more details about the tissue than the manual annotation. As an example, a large region annotated as ductal carcinoma *in situ* (DCIS) in patient H also contained spatial spots that were assigned to cluster H2, which corresponds to a region dense in invasive tumor cells, suggesting that this region has morphological properties of DCIS but molecular properties of invasive cancer (Fig. 2d). To further validate our clustering approach, we carried out differential gene expression analysis between each hierarchical cluster (Fig. 2a-c, Supplementary Fig. 4–6) and the rest of the tissue. A pathway analysis^29^ was then performed on the differentially expressed genes (Supplementary Table 3) to identify biological functions for various regions of the tumor sample (Supplementary Table 4). In most of the patients, we detected clusters in which the predominant pathways were clearly related to immune response. These clusters mainly overlapped with regions annotated as immune cells, tumor stroma/fibrous tissue and invasive cancer (Fig. 2e).

**Figure 2.**
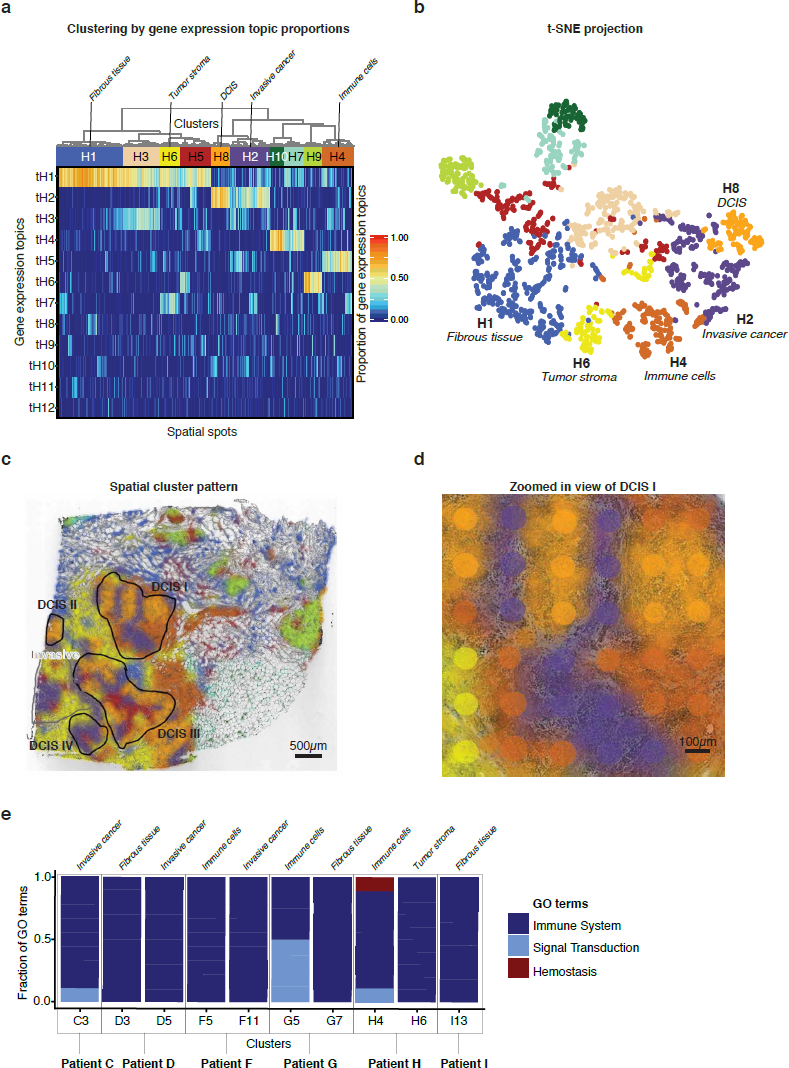
Clustering and pathway analysis. (a) The clustering was performed based on proportions of gene expression topics in patient H using Euclidean distance and Ward’s method. Colored bars under the dendrogram indicate cluster identity as determined by the unsupervised clustering method. Interesting clusters that clearly overlap with the manual annotations are marked. (b) Visualization of the clusters in t-SNE space. Data points represent spatial spots from patient H. Colors represents the cluster membership of each spatial spot. (c) Interpolated view across the tissue image for patient H. The different clusters show specific spatial patterns that closely follow the morphology. Black and gray lines bound areas determined to be DCIS and invasive cancer regions by the pathologist. The black bar in the bottom left corner represents 500µm. Colors represents the cluster membership of each spatial spot. (d) Magnified view of a DCIS area. The interpolated visualization is overlaid onto the actual spatial spots. The annotated DCIS region clearly contains spatial spots that are grouped into different clusters; for example, cluster H2, which dominates the invasive region. The black bar in the bottom left corner represents 100µm. (e) Pathway analysis of ten clusters across tumors from six of the patients. The pathway analysis is based on genes in the specific clusters that are upregulated relative to the rest of the tissue and show a high proportion of immune-related pathways (GO terms). Most immune-related GO terms were detected in immune cell, tumor stroma/fibrous tissue and invasive cancer regions.

**Figure 3.**
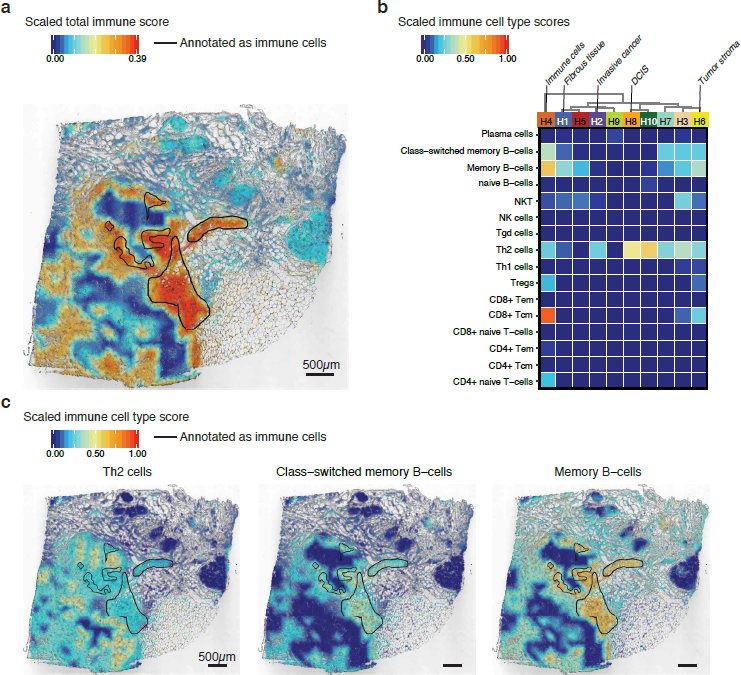
Spatial immune cell analysis. (a) A spatial view of the total immune score as calculated by xCell. The highest score was detected in a region annotated as immune cell dense (black lines). However, other regions also indicate high presence of immune cells, especially the tumor stroma (proximal to the tumor areas). The maximum score in the image is based on the patient max (0.39) after the score across all ten patients was scaled (0-1). The black bar in the bottom left corner represents 500µm. (b) Heat map of xCell scores for different immune cell-types across the ten clusters detected in patient H. Interesting clusters that clearly overlap with the manual annotation are marked. Immune cell cluster (H4) shows the highest score and immune cell diversity. However, several immune cell-types are present in other parts of the tissue. (c) A spatial view of the detected immune cell-types. Th2 cells are mostly present in the tumor regions but were also detected in the immune cell dense and stromal regions. Both Class-switched memory B-cells and memory B-cells are prevalent in the immune cell dense regions, but also appear in different parts of the stroma. The immune cell dense regions are marked with black lines in the images. The black bar in the bottom left corner of each tissue section represents 500µm.

**Figure 4.**
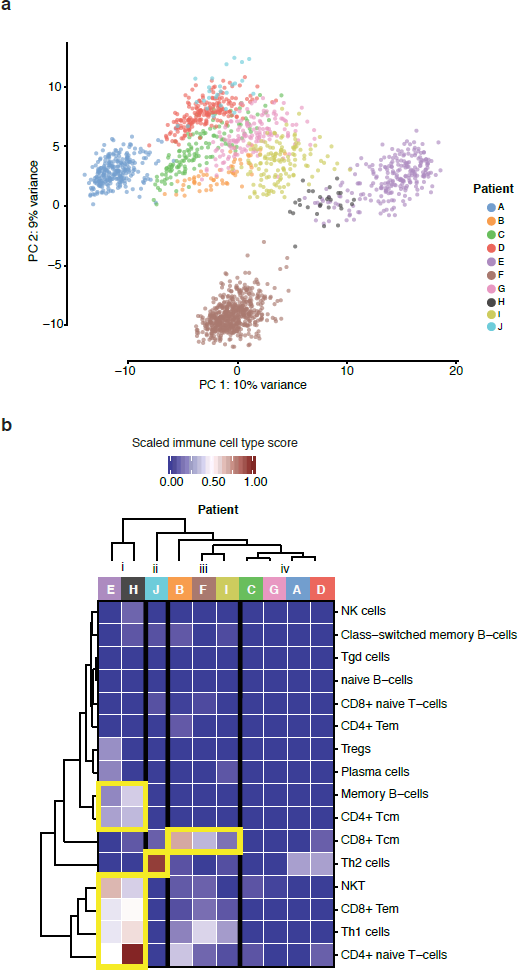
Lymphoid cell-types detected in the invasive regions across ten patients. (a) A PCA of spatial spots that were manually selected from the annotated invasive regions shows the formation of four major groups, with patients A and F separated from each other as well as from the other patients. Patients E and H formed their own group. The variance of each component is shown on the axes. (b) The xCell score was used to cluster the samples, and four clear clusters were detected (i-iv). Samples from patients E and H contained mostly NKT, CD8+ Tem, Th1 cells and CD4+ naïve T-cells, with some presence of Memory B-cells and CD4+ Tcm. Samples from patient J contained almost exclusively Th2 cells. Samples from patients B, F and I contained mostly CD8+ Tcm with the presence of some Th1 cells. Samples from patients C, G, A and D contained very few lymphoid cells, indicating lack of immune cell infiltration.

**Figure 5.**
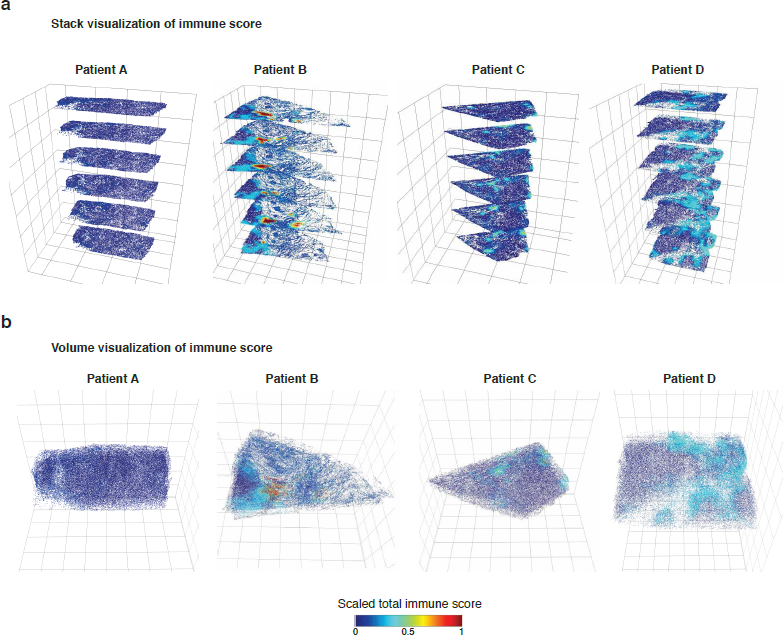
Three-dimensional Spatial Transcriptomics. Total immune score visualized in two different ways across six tissue sections from three patients. (a) Stacked view in which the sections are clearly separated for easy interpretation of expression patterns. (b) volume visualization, which is a more accurate view of the tissue volume. Clear differences in the spatial locations of regions with high immune scores can be noticed across all three dimensions. Patient B shows the highest immune score.

To further explore the data, we calculated spatial immune scores and determined the immune cell composition within each sample. The previously derived clusters served as the input for immune cell analysis with xCell^30^, which requires decent read depth regarding gene detection and counts to work properly. To obtain additional coverage and to overcome the exclusion of genes with low expression, we collapsed spatial spots within each cluster. This approach enabled us to detect almost six times more unique genes per sample (Supplementary Fig. 6c). The immune cell analysis revealed a high total immune score (the sum of several cell-types) for the annotated immune cell areas in a majority of the samples (Fig. 3a, Supplementary Fig. 7). Additionally, in several samples the total immune score overlapped well with the pathway analysis results (Fig. 2e). This approach also provided the immune cell composition for each of the clusters (Fig. 3b, Supplementary Fig. 8). For most samples, the highest values were detected in the regions annotated as immune cell dense. However, the scores for certain immune cell-types were relatively high even in other regions. For example, in patient H, Th2 (T helper cell 2), class-switched memory B-cells and memory B-cells were detected in the DCIS, tumor stroma, and all stroma regions, respectively (Fig. 3c). The predominant immune cell type identified in DCIS clusters from four patients (patient A: A13, patient G: G11, patient H: H8, patient J: J1 and J8) was Th2, a finding that suggests a homogeneous immune cell pattern within DCIS tissue (Fig. 3b, Supplementary Fig. 8).

Subsequent analysis focused on the immune cell composition of tumor regions annotated as invasive cancer across all ten patients. We selected spatial spots from the invasive regions of all tumor sections. A dimensionality reduction by PCA revealed four major groups, with patients A and F clearly separated from each other as well as the other patients. Patients E and H formed their own group (Fig. 4a), a finding that was also noted in the “bulk” analysis (Supplementary Fig. 1b). Furthermore, we used invasive spatial spots as the input for the immune cell analysis. The patient samples could be categorized into four groups (i-iv) based on immune cell composition (Fig. 4b). As was observed in the PCA, patients E and H also clustered together in this analysis, which indicates that the tumors of these two patients share a tumor infiltrating immune cell composition. The invasive cancer regions in samples from these two patients showed substantial immune cell diversity, containing mostly Natural killer cells (NKT), CD8+ Tem (effector memory T-cells), Th1 cells (T helper cell 1) and CD4+ naïve T-cells with some presence of Memory B-cells and CD4+ Tcm (central memory T-cells). Th1 cells are derived from activated CD4+ naïve T-cells, and can further activate CD8+ T-cells^31^. The presence of these types of tumor infiltrating CD8+ T-cells is correlated with better overall survival^12^–14. Moreover, NKTs have been shown to be highly cytotoxic to cancer cells and likely contributed to previously observed reductions in tumor cells^32,33^. The tumor sample from Patient J almost exclusively contained Th2 cells. As these cells are mainly responsible for recruiting and activating other immune cells, it is surprising that only this type of immune cell was detected. This finding might reflect failure to activate or attract other immune cell-types, which could negatively influence the overall survival in patients with similar immune cell composition. Tumor samples from three patients, B, F and I, showed mainly CD8+ Tcm (central memory T-cells) with some presence of Th1 cells. The immune cell pattern in these patients is similar to what was noted for patients E and H, but with less immune cell-type diversity. Tumor regions annotated as invasive cancer from the remaining four patients, C, G, A and D, showed a general absence of lymphoid cell signatures, indicating lack of immune cell infiltration. Patients with this type of pattern usually have poor overall survival^8,9^ and we observe that patients C and D are the only two deceased among the investigated patients (Supplementary Table 1).

Our analyses revealed high two-dimensional spatial variation in both gene expression and immune cell composition, which prompted a three-dimensional investigation of the patient samples. We selected samples from four of the patients and used cross-sectioning to collect six sections from each sample. The tissue image alignment provided a transformation matrix that we applied to the spatial spots and generated interpolated expression images into three-dimensional space (Supplementary Fig. 9). The output is a projection of the tissue in which any gene expression value, gene expression topic proportion or immune cell composition can be visualized. We applied the immune scores to the 3D reconstructed data and visualized the output in stacked images of each section (Fig. 5a) or by volume-based images (Fig. 5b). Patient A showed virtually no immune score whilst Patient B had the highest immune score among the four patients, but the high immune score was restricted to a few specific continuous nodes in space. Three of the patients showed variation across the additional dimension provided by this analysis, which demonstrates the importance of expanding the analysis from 2D to 3D. To further explore the 3D-data, we developed a R-shiny app (https://spatialtranscriptomics3d.shinyapps.io/ST3D-Viewer/) where all immune-cells and detected genes can be interactively visualized.

## Discussion

Here we investigate tumor heterogeneity in samples from ten patients diagnosed with HER2+ breast cancer in two and three dimensions using spatially resolved transcriptomics. We used novel methods to cluster the transcriptome wide data according to underlying structures in the form of LDA-based gene expression topics rather than pure gene expression values or morphological characteristics. By collapsing spatial spots within the clustering approach, we detected almost six times more genes than when using single spatial spots as the input for cell type analysis, improving the overall sensitivity of the downstream analysis. In most samples, our analysis revealed a clear overlap between the identified LDA clusters and manually annotated regions.

A key property of Spatial Transcriptomics technology is the possibility to identify and characterize the distribution of immune cells within tumor tissue sections in an unbiased way. Here, we applied spatial cell scoring to reveal areas with a presence of immune cells with the aim to study the immune composition within areas of invasive cancer of particular interest for survival outcome and immune therapy. We observe an array of distinct infiltrating immune cell patterns across the different patients. Although more extensive studies will be required to fully grasp the immune landscape, we observe groups of patients sharing similar immune repertoires in situ as well as patients lacking measurable immune cell infiltration. In the extension of this work it is likely that HER2+ tumors can be described not only by the gene expression profile per se but also by scoring the infiltrating immune cell profiles.

More extensive patient sampling is likely to establish a relationship between the extent of immune cell infiltration and node positivity. An interesting observation is that the two deceased patients belonged to the immune score group that showed a lack of immune cell infiltration in the invasive cancer region. Furthermore, tumors from two of the patients also showed the presence of CD4+ naïve T-cells, which are usually not present in tumor tissue^34^ but have earlier been reported in breast cancer samples^35^. Importantly, here we demonstrate that the described methodological approach can be used to dissect many different aspects of the immune-tumor interplay and substantiate whether the lack of immune cell infiltration or type of immune cells in tumor tissue is related to worse prognosis and/or lower overall survival.

This is the first attempt, to our knowledge, to present genome-wide RNA-seq data from human tissue in three dimensions. This is achieved by combining cross-sectioning with computational image alignment and data transformation. Our model demonstrated that it is possible to detect and follow immune score changes across all three dimensions, and that the analysis of a single, two-dimensional section would leave out important information regarding immune cell distribution within heterogeneous samples.

The resulting comprehensive view of gene expression in a tissue volume can thus facilitate new understanding of tumors and the surrounding microenvironment and has the potential to challenge current diagnostic practices. By nature, cancers are heterogeneous, and the presented results increase the notion of complexity to a higher degree. Yet, the analysis demonstrates that using a three-dimensional description of the tumor tissue landscape can advance diagnostic procedures and help design personalized treatments from the time of diagnosis.

### Acknowledgements

We thank Janne Lehtiö, Therese Sørlie, Sten Linnarsson, Fredrik Pontén, David Redin, Phil Ewels and Lennart Kester for valuable contributions. The data were analyzed using SNIC and Uppsala Multidisciplinary Center for Advanced Computational Science (SNIC/UPPMAX) resources. This work was supported by the Knut and Alice Wallenberg Foundation, Swedish Cancer Society, Swedish Foundation for Strategic Research, the Swedish Research Council, Tobias Stiftelsen, Torsten Söderbergs Foundation and Science for Life Laboratory. We thank the National Genomics Infrastructure (NGI), Sweden for providing infrastructural support. The raw data were deposited at the NCBI database of Genotypes and Phenotypes (dbGAP) with accession number PRJNA354717. Images, gene counts and scripts can be accessed at www.spatialtranscriptomicsresearch.org.

## Author Contributions

F. S. and P. L. S. wrote the manuscript with input from all other authors. L. S. carried out the laboratory experiments. L. L., F. S. and S. V. performed data analysis with input from J. V. C. and J. H. A. E. inspected the patient samples and performed morphological annotations. Å. B., J. V. C. and J. H. L. S. provided samples and biological input. J. L., P. L. S., J. F. and Å. B. planned the study.

## Methods

### Array production

The array production was described previously^22,36^. Briefly, the microarrays were generated as a 33×35 matrix with a 200 µm center-to-center distance between 100 µm spatial spots. A total of 1007, unique and spatially barcoded DNA oligonucleotides, were used.

### Tissue handling, staining and imaging

These steps were described previously^22^. Shortly, fresh frozen material was sectioned at 16 µm. After placing the tissue on top of the barcoded microarray, the glass slide was warmed at 37 °C for 1 min for tissue attachment and fixated in ˜ 4% NBF (neutral buffered formalin) for 10 min at room temperature (RT). The slide was then washed briefly with 1x PBS (phosphate buffered saline). The tissue was dried with isopropanol before staining. The tissue was stained with Mayer’s hematoxylin for 4 min, washed in Milli-Q water, incubated in bluing buffer for 2 min, washed in Milli-Q water, and further incubated for 1 min in 1:20 eosin solution in Tris-buffer (pH 6). The tissue sections were dried for 5 min at 37 °C and then mounted with 85% glycerol and a coverslip. Imaging was performed using the Metafer VSlide system at 20x resolution. The images were processed with the VSlide software (v1.0.0). After the imaging was complete, the cover slip and remaining glycerol were removed by dipping the whole slide in Milli-Q water followed by a brief wash in 80% ethanol and warming for 1 min at 37 °C.

### Permeabilization and cDNA synthesis

Permeabilization and cDNA synthesis were carried out as previously described^22^ but with substituting the Exonuclease I buffer pre-permeabilization treatment with a 20 min incubation at 37 °C in 14 U of collagenase type I (Life Technologies, Paisley, UK) diluted in 1x HBSS buffer (Thermo Fisher Scientific, Life Technologies, Paisley, UK) supplemented with 14 µg BSA followed by an incubation in 0.1% pepsin-HCl (pH 1) for 10 min at 37 °C. A cDNA-mix containing Superscript III, RNaseOUT, DTT, dNTPs, BSA and Actinomycin D was added and the slide incubated at 42 °C overnight (˜18 h). The tissue was washed with 0.1x SSC between each incubation step.

### Tissue removal and cDNA release from surface

Tissue removal, as well as the release of cDNAs from surface was described previously^22^. In brief, beta-Mercaptoethanol was diluted in RNeasy lysis buffer and samples were incubated for 1 h at 56 °C. The wells were washed with 0.1x SSC followed by incubation with proteinase K, diluted in proteinase K digestion buffer, for 1 h at 56 °C. The slides were then washed in 2x SSC + 0.1% SDS, 0.2x SSC followed by 0.1x SSC and dried. The release mix consisted of second strand buffer, dNTPs, BSA and USER enzyme and was carried out for 2 h at 37 °C. After probe release, the 1007 spatial spots containing non-released DNA oligonucleotide fragments were detected by hybridization and imaging, in order to obtain Cy3-images for alignment.

### Library preparation and sequencing

The protocol followed the same preparation procedures as described earlier^22^, but were carried out using an automated pipetting system (MBS Magnatrix Workstation), also previously reported^27^. In general, second strand synthesis and blunting were carried out by adding DNA polymerase I, RNase H and T4 DNA polymerase. The libraries were purified and amplified RNA (aRNA) was generated by a 14 h *in vitro* transcription (IVT) reaction using T7 RNA polymerase, supplemented with NTPs and SUPERaseIN. The material was purified and an adapter ligated to the 3’-end using a truncated RNA ligase 2. Generation of cDNA was carried out at 50 °C for 1 h by Superscript III, supplemented with a primer, RNaseOUT, DTT and dNTPs. Double stranded cDNA was purified, and full Illumina sequencing adapters and indexes were added by PCR using 2xKAPA HotStart ready-mix. The number of amplification cycles needed for each section was determined by qPCR with the addition of EVA Green. Final libraries were purified and validated using an Agilent Bioanalyzer and Qubit before sequencing on the NextSeq500 (v2) at a depth of ˜100 million paired-end reads per tissue section. The forward read contained 31 nucleotides and the reverse read 46 nucleotides.

### Mapping, gene counting and demultiplexing

These steps were carried out in a similar fashion to what has previously been described^22^. The forward read contained the spatial barcode and a semi-randomized UMI sequence (WSNNWSNNV) while the reverse read contained the transcript information and was used for mapping to the reference GRCh38 human genome. Before mapping the reads with STAR^37^, the reverse reads were first quality trimmed based on the Burrows-Wheeler aligner and long homopolymer stretches removed. HTSeq-count^38^ with the setting *-intersection-nonempty*, was used to count only protein-coding and long non-coding transcripts for each gene, using an Ensembl reference file (v.79). The remaining reads were taken into TagGD demultiplexing^39^ using the 18 nucleotides spatial barcode. The demultiplexed reads were then filtered for amplification duplicates using the UMI with a minimal hamming distance of 2. The UMI-filtered counts were used in the analysis. The analysis pipeline (v0.8.5) is available at https://github.com/SpatialTranscriptomicsResearch/st_pipeline.

### Analysis of bulk data

The data from all spatial spots, for each tumor or section, were added up separately. The data were either used of PCA or AIMS^40^ to determine subtypes.

### LDA and clustering

Replicate datasets from each tissue were merged and a Latent Dirichlet Allocation (LDA) model for each merged dataset using the R package cellTree^25^. The numbers of topics for each dataset were chosen based the Bayes factor over the Null model^41^ using the “maptpx” method with a maximum number of allowed topics set to 15. The resulting topic matrices were used as a basis for hierarchical clustering of spatial spots. Clusters were chosen using the adaptive method Dynamic Tree Cut^42^. Clustered spatial spots were color coded and visualized on a heatmap of topic/spatial spot pairs and overlaid on the tissue images. t-SNE based on topic proportions was computed for each sample with each point colored by its respective cluster identity.

### Cell-type enrichment analysis

Each cluster was collapsed into vectors by adding the gene expression values for each spatial spot within that cluster. Cell-type enrichment was performed for each cluster using the gene signature based method xCell^30^ or each tissue sample. xCell scores for cluster/cell-type pairs were visualized as heatmaps. Cluster/cell-type pair scores were then extrapolated to the spatial spots within each respective cluster to generate matrices with xCell scores for spatial spot/cell-type pairs.

### Visualization of cell distribution and interpolation

Tissue images in jpeg format were converted to grey scale and point scatters of x, y coordinates were generated by defining points at pixel coordinates with intensity below a threshold of 0.5. The 2D point scatters were transformed from pixel coordinates to array coordinates, thus defined in the same coordinate system as the array spatial spots. Next, a raster was generated across each tissue image and points in the 2D scatter were associated with a grid cell by calculating the minimum Euclidean distance. Spatial spot topic proportions and xCell scores were interpolated^43^ across the raster and assigned to each point of the 2D scatter. The 2D scatters were overlaid onto the tissue images and colored by either topic proportions or xCell scores. A subset of xCell scores was selected to include only immune cells and was scaled across all immune cell-types and replicates to range between 0 and 1. The immune cell-type group included; CD4+ memory T-cells, CD4+ naïve T-cells, CD4+ Tcm, CD4+ Tem, CD8+ naive T-cells, CD8+ Tcm, CD8+ Tem, Tregs, Th1 cells, Th2 cells, Tgd cells, NK cells, NKT, naïve B-cells, Memory B-cells, Class-switched memory B-cells, pro B-cells and Plasma cells. For generation of 3D data, all respective sections for each sample were processed with Fiji using the “Transform Virtual Stack Slices” plugin^44^. This created a transformation matrix for each registered image and enabled image alignment. The transformation matrix was used to transform^45^ the registered binary dots and spatial spot coordinates. The transformed dots and spatial spots were scaled, centered, manually inspected and re-adjusted if necessary. Each tissue section was multiplied three times and stacked by separating each section with an even distance along the z-axis. xCell scores were generated for clusters spanning the whole 3D volume and the values were interpolated onto the 2D scatters separately. All the stacks from each section and sample were combined and visualized as interactive 3D scatter heatmaps (HTMLwidgets) using the R package plotly^46^. xCell scores were scaled to range between 0 and 1 on the color scale.

### DE and pathway analysis of clusters

Filtered gene expression data was normalized as counts per ten thousand by dividing each spatial spot column by its sum of counts and multiplying by 10,000. Spatial spots grouped by LDA based clusters were normalized using scran^47^. First, each cluster was filtered from spatial spots with 0 value size factors and clusters with less than 40 spatial spots were discarded. The remaining clusters were normalized with the computed size factors. DE analysis was performed using an edgeR workflow^48^. For each cluster, a design matrix was constructed grouping clustered spatial spots and all remaining spatial spots separately. Estimates of common and trended dispersions were computed using the “estimateDisp” function and negative binomial generalized log-linear models were fitted to each design matrix using the “glmFit” function. Likelihood ratio tests were calculated using the “glmLRT” function to obtain differentially expressed genes with a log2-fold change greater than 1 at a significance threshold of p = 0.01.

### Analysis of annotated regions

Spatial spots within regions annotated as “mainly invasive cancer”, “carcinoma *in situ*” and “inflammatory cells” were selected using the Spatial Transcriptomics Research Viewer and exported as expression matrices in tsv-format. Annotated groups were pooled and subjected to cell-type enrichment with xCell and the results were visualized as heatmaps of xCell scores for cluster/cell-type pairs. The Spatial Transcriptomics Research Viewer is available at https://github.com/SpatialTranscriptomicsResearch/st_viewer.

---

**Supplementary Fig. 1.**
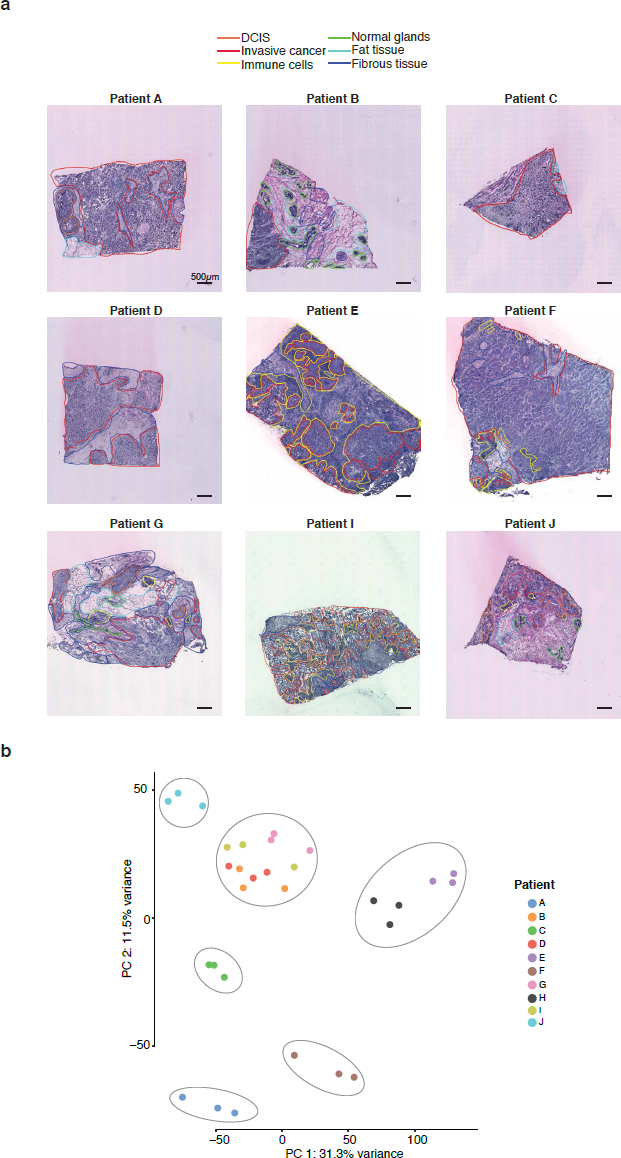
Overview of patient samples. (a) Morphological regions were characterized by a pathologist and annotated into six distinct categories: DCIS (orange); invasive cancer (red); immune cells (yellow); normal glands (green); fat tissue (cyan); and fibrous tissue (blue). The black bar in the bottom left corner of each tissue section represents 500µm. (b) PCA of bulk data from the ten patients shows that the patients can be categorized into separate groups, indicating large interpatient differences. However, patients B, D, G and I and patient E and H group together, indicating transcriptional similarities. The variance of each component is shown on the axes.

**Supplementary Fig. 2.**
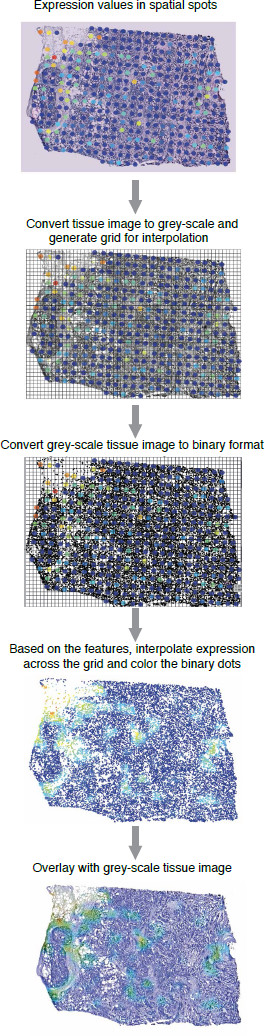
Schematic representation of the interpolation method. Spatial spot values, e.g. gene expression or proportions of gene expression topics, were overlaid with the HE-stained tissue image. HE-stained tissue images were first converted into gray-scale and then into binary format based on a light intensity threshold. Tissue images were then rasterized to generate a grid of cells into which the binary dots were binned according to the minimum Euclidean distance from each binary dot to the grid cell centers. Spatial spots values were interpolated across the grid and assigned to each grid-cell. The binary dot patterns were colored based on the interpolated values and overlaid on top of the gray-scale tissue images.

**Supplementary Fig. 3.**
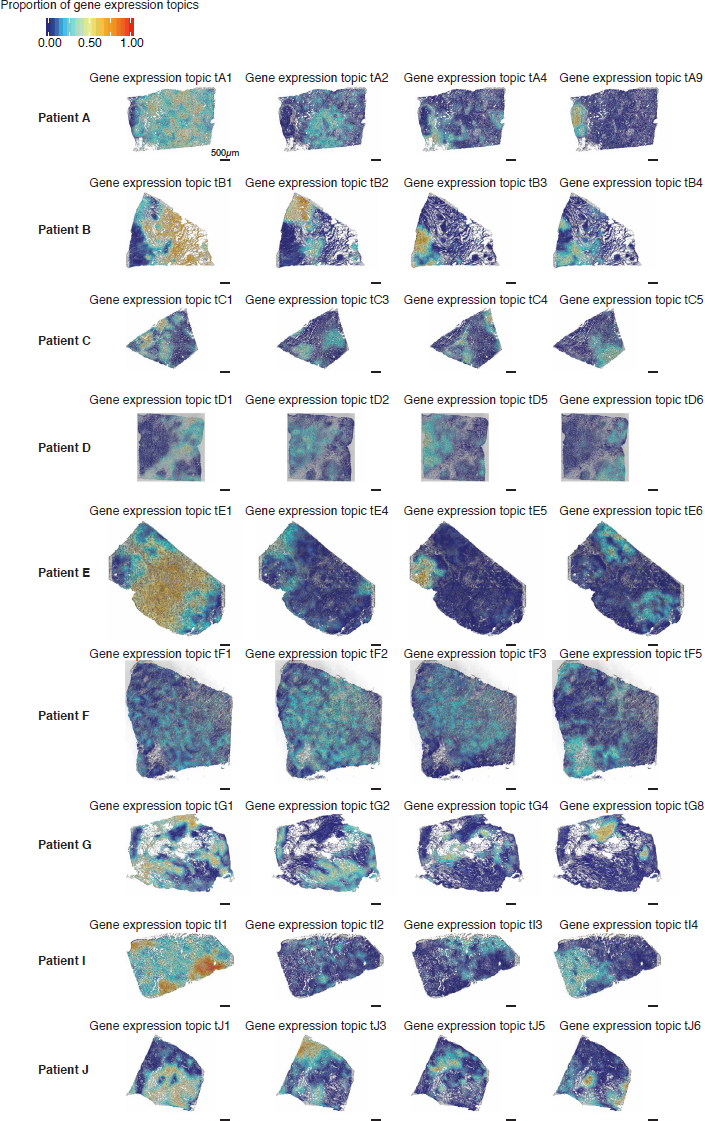
A spatial visualization of the proportions of gene expression topics. Interpolated tissue images show four gene expression topics with clear morphological patterns. Gene expression topic proportions are sample specific, meaning that the same proportion in another sample does not necessarily represent a similar region. The black bar in the bottom left corner of each tissue section represents 500µm.

**Supplementary Fig. 4.**
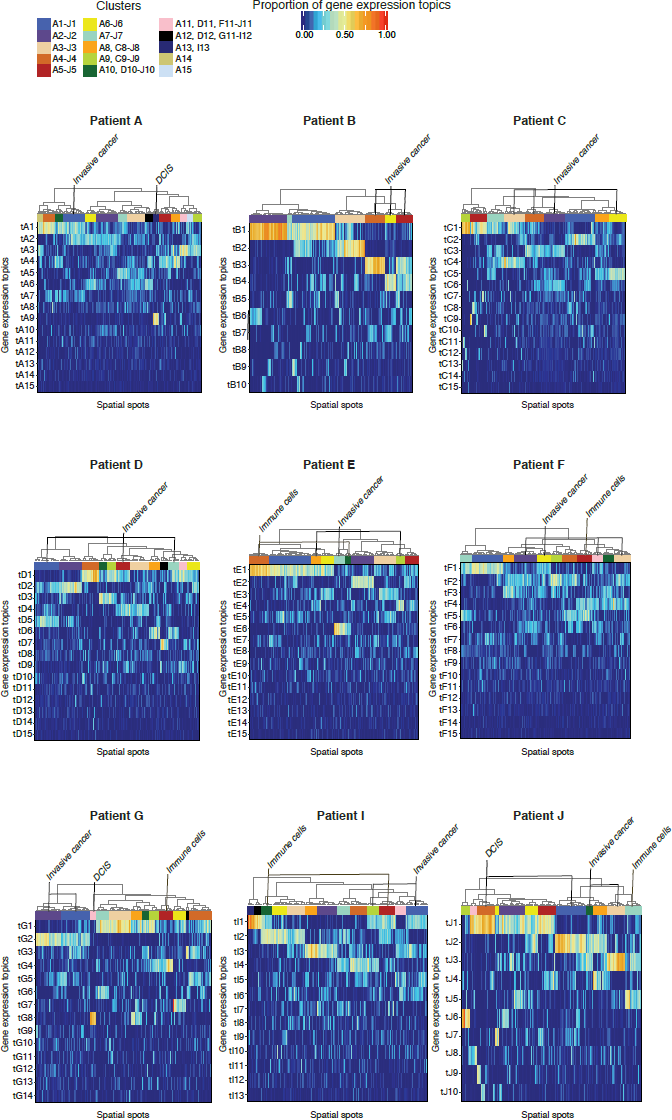
Hierarchical clustering of spatial spots. The clustering was based on proportions of gene expression topics in patient tissue samples and performed using Euclidean distance and Ward’s method. Reference bars under the dendrograms indicate cluster identity determined by the unsupervised clustering method. Interesting clusters that clearly overlap with the manual annotations are named. Clusters are sample specific, meaning that the same color does not represent a specific region across different samples.

**Supplementary Fig. 5.**
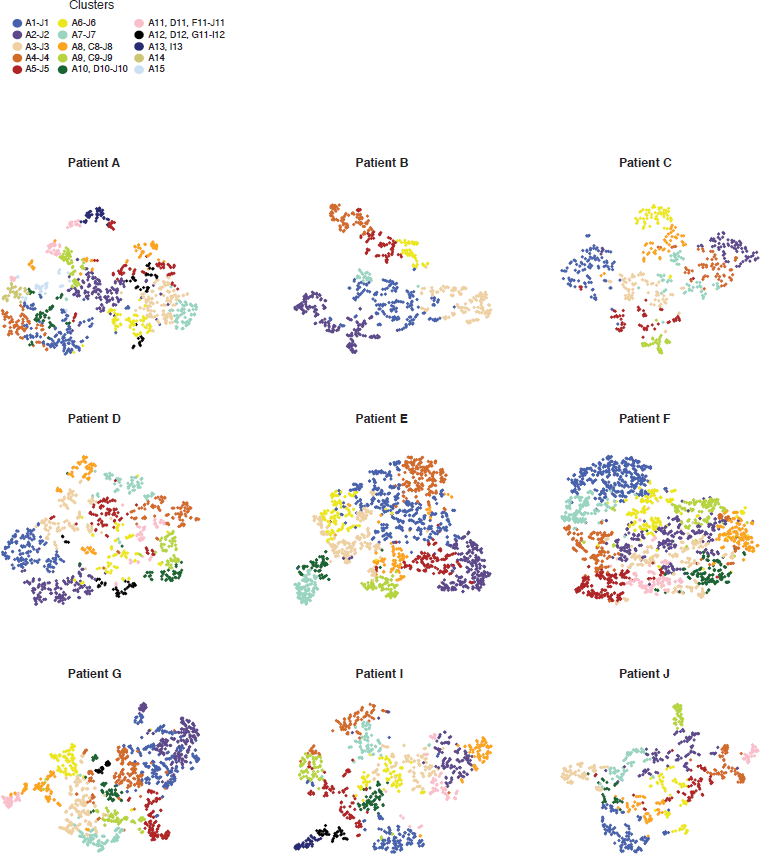
Non-linear dimensionality reduction using t-SNE. Data points represents spatial spots and are colored based on cluster identity. The plot was used to verify the clustering. Clusters are sample specific, meaning that the same color does not represent a specific region across different samples.

**Supplementary Fig. 6.**
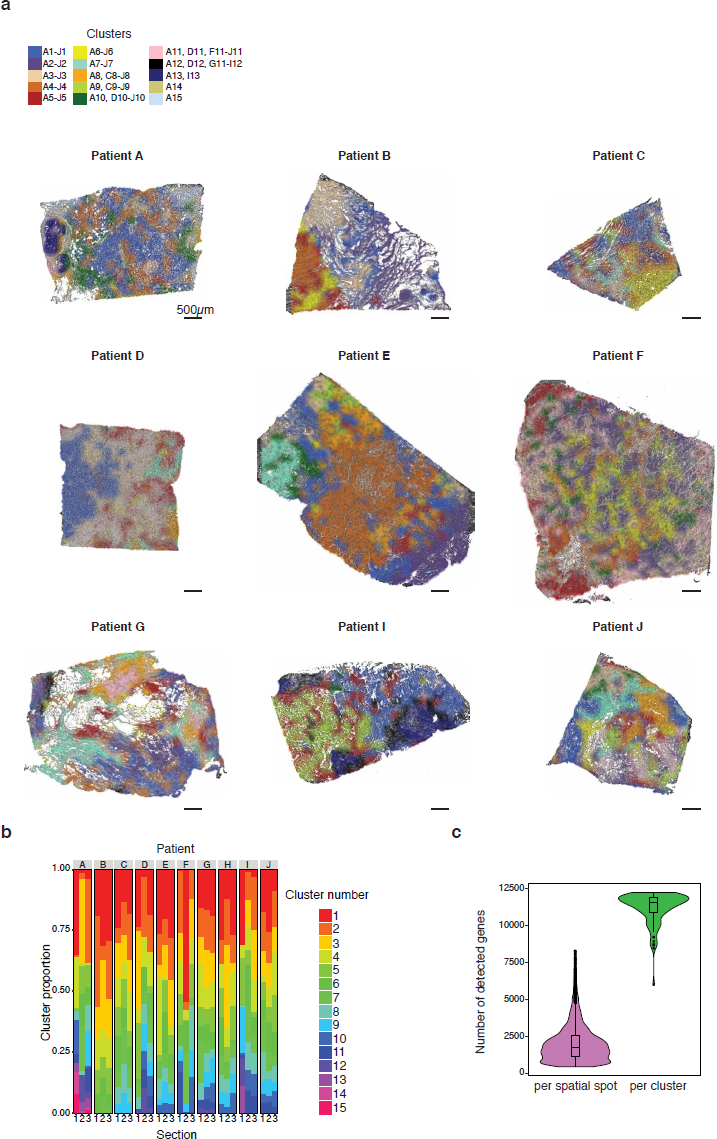
Spatial visualization of clusters. (a) Clusters were classified by color and interpolated across tissue images. Clusters are sample specific, meaning that the same color does not represent a specific region across different samples. The black bar in the bottom left corner of each tissue section represents 500µm. (b) Stacked bar chart of cluster proportions across all three sections (replicates) of each of the ten patient samples. Similar bar patterns indicate similar cluster proportions across replicates. (c) Violin plot of the number of unique genes that are detected per spatial spot in relation to how many are detected per cluster when spatial spots are collapsed.

**Supplementary Fig. 7.**
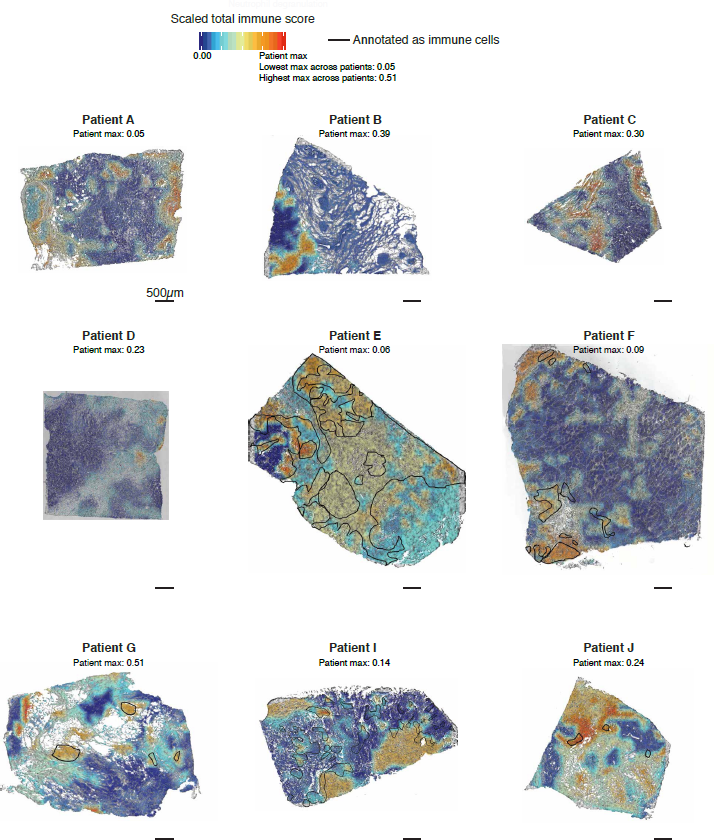
Spatial visualization of total xCell immune scores. Interpolated tissue images demonstrate how the immune score is distributed across the tumor tissue. The color scale (top left) shows the extent of immune cell density while the black lines display manually annotated immune cell dense regions. Maximum scores in the images are based on the patient max after the score across all ten patients were scaled (0-1). The black bar in the bottom left corner of each tissue section represents 500µm.

**Supplementary Fig. 8.**
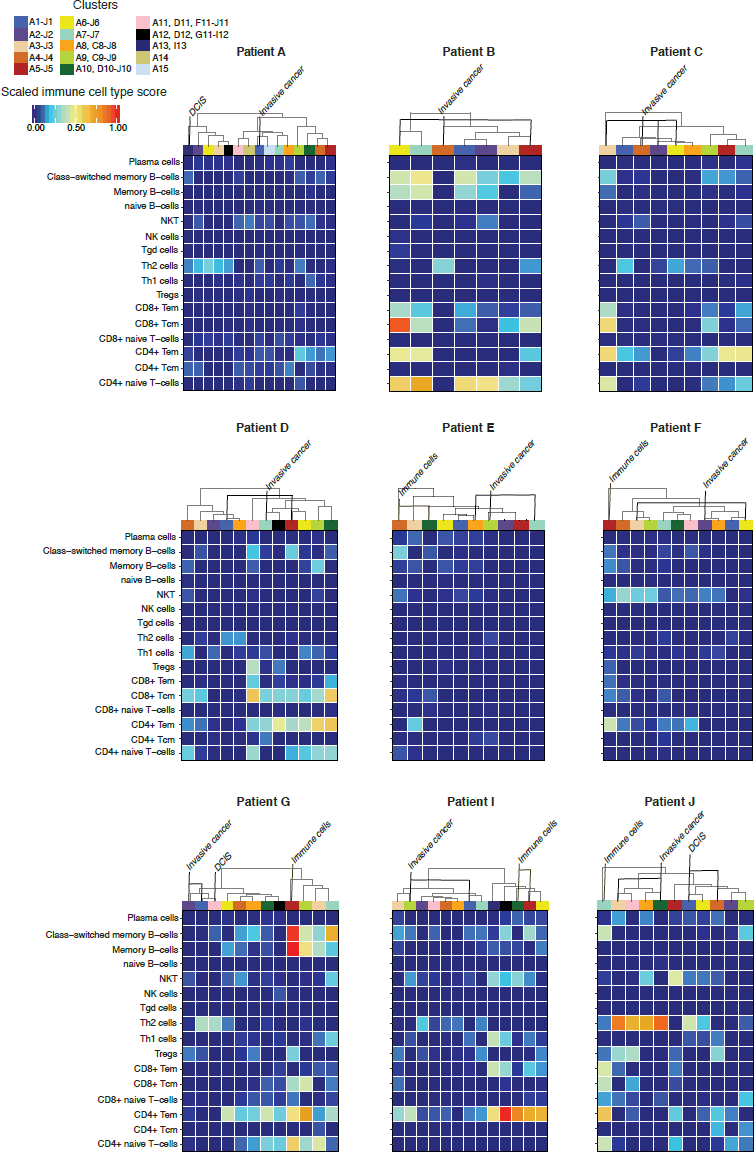
Heatmap representation of immune cell-type scores among clusters. The xCell scores were calculated for each cluster (columns) and patient sample (heatmaps) separately. Each grid-cell represents the xCell score for a certain lymphoid cell-type in a specific cluster. The values have been scaled across all samples to a range between 0 and 1. Interesting clusters that clearly overlap with the manual annotations are marked.

**Supplementary Fig. 9.**
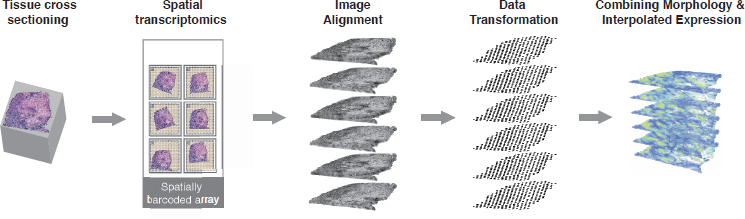
Three-dimensional Spatial Transcriptomics. Schematic representation of the approach. Six sections were collected from each of the four samples and processed using the Spatial Transcriptomics methods. Tissue images were computationally aligned and stacked. The transformation matrix from the alignment were applied to the spatial data so that the images and data existed in the same space. The data were then interpolated as previously described.

## References

1. López-otín C., Blasco M. A., Partridge L., Serrano M. & Kroemer G. Europe PMC Funders Group The Hallmarks of Aging. Cell 153, 1194–1217 (2013).

2. Magalhães J. P. De. How ageing processes influence cancer. 13, (2013).

3. Carstens J. L. et al. Spatial computation of intratumoral T cells correlates with survival of patients with pancreatic cancer. Nat. Commun. 8, 1–13 (2017).

4. Lyons Y. A., Wu S. Y., Overwijk W. W., Baggerly K. A. & Sood, A. K. Immune cell profiling in cancer: molecular approaches to cell-specific identification. npj Precis. Oncol. 1, 26 (2017).

5. Gajewski T. F., Schreiber H. & Fu, Y.-X. Innate and adaptive immune cells in the tumor microenvironment. Nat. Immunol. 14, 1014 (2013).

6. Zaha D. C. Significance of immunohistochemistry in breast cancer. World J. Clin. Oncol. 5, 382 (2014).

7. Prat A. et al. Molecular features and survival outcomes of the intrinsic subtypes within HER2-positive breast cancer. J. Natl. Cancer Inst. 106, 1–8 (2014).

8. Stanton S. E. & Disis M. L. Clinical significance of tumor-infiltrating lymphocytes in breast cancer. J. Immunother. Cancer 4, 59 (2016).

9. García-Teijido P., Cabal M. L., Fernández I. P. & Pérez Y. F. Tumor-infiltrating lymphocytes in triple negative breast cancer: The future of immune targeting. Clin. Med. Insights Oncol. 10, 31–39 (2016).

10. Khan A. M. & Yuan Y. Biopsy variability of lymphocytic infiltration in breast cancer subtypes and the ImmunoSkew score. Sci. Rep. 6, 2–11 (2016).

11. Degnim A. C. et al. Immune cell quantitation in normal breast tissue lobules with and without lobulitis. Breast Cancer Res. Treat. 144, 539–549 (2014).

12. Ali H. R. et al. Association between CD8+ T-cell infiltration and breast cancer survival in 12 439 patients. Ann. Oncol. 25, 1536–1543 (2014).

13. Mahmoud S. M. A. et al. Tumor-Infiltrating CD8+ Lymphocytes Predict Clinical Outcome in Breast Cancer. J. Clin. Oncol. 29, 1949–1955 (2011).

14. Miyashita M. et al. Prognostic significance of tumor-infiltrating CD8+ and FOXP3+ lymphocytes in residual tumors and alterations in these parameters after neoadjuvant chemotherapy in triple-negative breast cancer: a retrospective multicenter study. Breast Cancer Res. 17, 124 (2015).

15. Huang Y. et al. CD4+ and CD8+ T cells have opposing roles in breast cancer progression and outcome. Oncotarget 6, 17462–78 (2015).

16. Yang J., Li X., Liu X. P. & Liu Y. The role of tumor-associated macrophages in breast carcinoma invasion and metastasis. Int. J. Clin. Exp. Pathol. 8, 6656–6664 (2015).

17. Obeid E., Nanda R., Fu Y. X. & Olopade O. I. The role of tumor-associated macrophages in breast cancer progression (review). Int. J. Oncol. 43, 5–12 (2013).

18. Williams C. B., Yeh E. S. & Soloff A. C. Tumor-associated macrophages: unwitting accomplices in breast cancer malignancy. npj Breast Cancer 2, 15025 (2016).

19. Savas P. et al. Clinical relevance of host immunity in breast cancer: from TILs to the clinic. Nat. Rev. Clin. Oncol. 13, 228 (2015).

20. Schumacher T. N. & Schreiber R. D. Neoantigens in cancer immunotherapy. Science (80-.). 348, 69 LP-74 (2015).

21. Farkona S., Diamandis E. P. & Blasutig I. M. Cancer immunotherapy: The beginning of the end of cancer? BMC Med. 14, 1–18 (2016).

22. Ståhl P. L. et al. Visualization and analysis of gene expression in tissue sections by spatial transcriptomics. Science (80-.). 353, 78 LP-82 (2016).

23. Giacomello S. et al. Spatially resolved transcriptome profiling in model plant species. Nat. Plants 3, 17061 (2017).

24. Blei D. M., Ng A. Y. & Jordan M. I. Latent Dirichlet Allocation. 3, 993–1022 (2003).

25. duVerle D. A., Yotsukura S., Nomura S., Aburatani H. & Tsuda K. CellTree: An R/bioconductor package to infer the hierarchical structure of cell populations from single-cell RNA-seq data. BMC Bioinformatics 17, 1–17 (2016).

26. Dey K. K., Hsiao C. J. & Stephens M. Visualizing the structure of RNA-seq expression data using grade of membership models. PLOS Genet. 13, e1006599 (2017).

27. Jemt A. et al. An automated approach to prepare tissue-derived spatially barcoded RNA-sequencing libraries. Sci. Rep. 6, 37137 (2016).

28. Asp M. et al. Spatial detection of fetal marker genes expressed at low level in adult human heart tissue. Sci. Rep. 7, 12941 (2017).

29. Yu G. & He Q.-Y. ReactomePA: an R/Bioconductor package for reactome pathway analysis and visualization. Mol. Biosyst. 12, 477–479 (2016).

30. Aran D., Hu Z. & Butte A. J. xCell: Digitally portraying the tissue cellular heterogeneity landscape. Genome Biol. 18, (2017).

31. Ekkens M. J. et al. Th1 and Th2 cells help CD8 T-cell responses. Infect. Immun. 75, 2291–2296 (2007).

32. Ames E., Hallett W. H. D. & Murphy W. J. Sensitization of human breast cancer cells to natural killer cell-mediated cytotoxicity by proteasome inhibition. Clin. Exp. Immunol. 155, 504–513 (2009).

33. Tallerico R. et al. NK cells control breast cancer and related cancer stem cell hematological spread. Oncoimmunology 6, e1284718 (2017).

34. Luckheeram R. V., Zhou R., Verma A. D. & Xia B. CD4+T cells: Differentiation and functions. Clin. Dev. Immunol. 2012, (2012).

35. Su S. et al. Blocking the recruitment of naive CD4+ T cells reverses immunosuppression in breast cancer. Cell Res. 27, 461–482 (2017).

36. Vickovic S. et al. Massive and parallel expression profiling using microarrayed single-cell sequencing − Accepted. Nat. Commun. 1–9 (2016). doi:10.1038/ncomms13182

37. Dobin A. et al. STAR: ultrafast universal RNA-seq aligner. Bioinformatics 29, 15–21 (2013).

38. Anders S., Pyl P. T. & Huber W. HTSeq-A Python framework to work with high-throughput sequencing data. Bioinformatics 31, 166–169 (2015).

39. Costea P. I., Lundeberg J. & Akan P. TagGD: Fast and Accurate Software for DNA Tag Generation and Demultiplexing. PLoS One 8, e57521 (2013).

40. Paquet E. R. & Hallett M. T. Absolute assignment of breast cancer intrinsic molecular subtype. J. Natl. Cancer Inst. 107, 1–9 (2015).

41. Taddy M. a. On Estimation and Selection for Topic Models. Proc. Fifteenth Int. Conf. Artif. Intell. Stat. (AISTATS 2012) 1184–1193 (2012).

42. Langfelder P., Zhang B. & Horvath S. Defining clusters from a hierarchical cluster tree: The Dynamic Tree Cut package for R. Bioinformatics 24, 719–720 (2008).

43. Akima H. A Method of Bivariate Interpolation and Smooth Surface Fitting for Irregularly Distributed Data Points. ACM Trans. Math. Softw. 4, 148–159 (1978).

44. Schindelin J. et al. Fiji − an Open Source platform for biological image analysis. Nat. Methods 9, 10.1038/nmeth.2019 (2012).

45. Carrillo G. vec2dtransf: 2D Cartesian Coordinate Transformation. (2015).

46. Sievert C. et al. plotly: Create Interactive Web Graphics via ‘plotly.js’. (2016).

47. L. Lun, A. T. Bach, K. & Marioni, J. C. Pooling across cells to normalize single-cell RNA sequencing data with many zero counts. Genome Biol. 17, 75 (2016).

48. Robinson M. D., McCarthy D. J. & Smyth G. K. edgeR: A Bioconductor package for differential expression analysis of digital gene expression data. Bioinformatics 26, 139–140 (2009).

